# Leaf susceptibility of Macaronesian laurel forest species to *Phytophthora ramorum*

**DOI:** 10.1101/2023.07.15.549153

**Authors:** Eduardo Moralejo, José A. García-Muñoz, Sandra Denman, Àlex Giménez-Romero

## Abstract

*Phytophthora ramorum* (*Pr*) is an invasive oomycete in Europe and North America and the causal agent of sudden oak death (SOD), which occurs along the coastal fog belt of California and southwestern Oregon, and it also causes sudden larch death in the UK. The Macaronesian laurel forest (MLF), a relict subtropical evergreen forest of the North Atlantic islands, shares climatic and some taxonomic affinities with those areas affected by SOD. To assess the disease risk, we tested the foliage susceptibility of MLF species and their capacity to sustain *Pr* sporulation and compared the climatic suitability with other areas where the pathogen is established. Detached leaves of 15 species were inoculated with zoospores and mycelium (through wounding) with five *Pr* isolates belonging to the EU1 and NA1 clonal lineages. MLF species showed diverse responses to *Pr*, ranging from extensive necroses on *Viburnum tinus* to asymptomatic sporulation on *Picconia excelsa*. Eleven species developed necrotic lesions to different degrees through zoospore inoculation while this increased to 13 species through wound treatment. Overall, small necrotic lesions (i.e. tolerance) were predominant, but *Pr* was rather aggressive to *V. tinus*, *Arbutus canariensis* and *Ilex canariensis*. Although the mean sporangial production was generally low (25-201 sporangia) in all species, the number of sporangia per leaf in five MLF species was similar to those reported for *Umbellularia californica*, a key host driving the SOD epidemics in California. Climatic suitability indexes in MLF areas were similar to those where SOD is found in California. Our results indicate a moderate to high risk of *Pr* establishment if the pathogen is introduced in the MLF.

## Introduction

The oomycete *Phytophthora ramorum* (*Pr*) is a superb example of a multiple-plant host pathogen (Grünwald et al., 2012). It causes sudden oak death (SOD), occurring in California and Oregon since the mid-1990s (Rizzo et al., 2002; Hansen et al., 2003), and sudden larch death in the UK, Ireland and France since 2009 (Brasier & Webber 2010; O’Hanlon et al., 2018; Schenck et al., 2018). *Pr* exhibits a wide host range in natural ecosystems and in ornamental plants in nurseries in Europe and North America (Werres et al., 2001; Rizzo et al., 2002; Grünwald et al., 2019). To date, this oomycete is known to infect leaves, twigs and stems of many unrelated taxa (e.g. ferns, angiosperms and gymnosperms) under mesic to humid conditions in Mediterranean and temperate-oceanic climates (Brasier et al., 2004; Davidson et al., 2005; Denman et al., 2006). In laboratory inoculations, *Pr* potential hosts include many ornamental plants (Tooley et al., 2004; Garbelotto et al., 2020) and wild trees from the east and west coasts of the USA (Hansen et al., 2005; Tooley & Browning 2009), New Zealand (Hüberli et al., 2008), Australia (Ireland et al., 2012) and Europe (Denman et al., 2005; Harris et al.,2021), including the Mediterranean basin (Moralejo et al., 2006b; Moralejo et al., 2009).

Understanding *Pr* pathogenic and fitness traits conferring the ability to invade plant communities may be useful for predicting disease risk for other ecosystems not yet exposed to the pathogen (Harris et al., 2021). Epidemiological studies of SOD in natural environments in California reveal that *Pr* is mainly an oak-laurel forest disease in which bay laurel (*Umbellularia californica*) and tanoak (*Notholithocarpus densiflorus*) act as key hosts driving epidemics (Davidson et al., 2005; Maloney et al., 2005; Anacker et al., 2008; Davidson et al., 2008; Meentemeyer et al., 2008; Kozanitas et al., 2022). Bay laurel often supports many infected leaves with tip and margin necrosis, which sustain *Pr* sporangial production, apparently without causing debilitating damage to the host, i.e. tolerance (Di Leo et al., 2009). By contrast, the stems of susceptible oaks adjacent to infected bay laurels may become infected from rain-splashed inoculum emanating from the bay laurel leaves and infections usually develop into lethal live bark cankers (Davidson et al., 2005).

Oak-laurel forests are the dominant vegetation type in tropical and subtropical Asian mountains, from the Himalayas to the Indomalayan archipelago, and support vegetation closely related taxonomically to the temperate evergreen oak forests of East Asia (Tagawa 1995). Recently, *Pr* has been discovered in the mountain laurel-oak forests of northern Vietnam and southwest Japan (Jung et al., 2021), confirming previous speculations of its centre of origin in east Asia (Rizzo et al., 2002; Brasier 2003; Kluza et al., 2007; Brasier et al., 2010). Oak-laurel and redwood forests also thrive along the fog belt of the Pacific coast of the USA, characterised by a Mediterranean climate with narrow seasonal and moderate daily temperature fluctuations and a long drought period in summer (Johnstone & Dawson 2010).

This type of climate has strong affinities with those where the Macaronesian laurel forest (MLF) has survived in the Canary Islands, Madeira and Azores (Figure 1). MLF represents the relict of the Neogene oak-laurel forest that once extended throughout southern and central Europe (Guimaräes & Olmeda 2008). Today, MLF is restricted to the cloud belt situated between 500 to 1500 m altitude on the northern slopes of the islands (15-40° latitude) under the direct influence of the northeastern trade winds (Guimaräes & Olmeda 2008).

**Figure 1.**
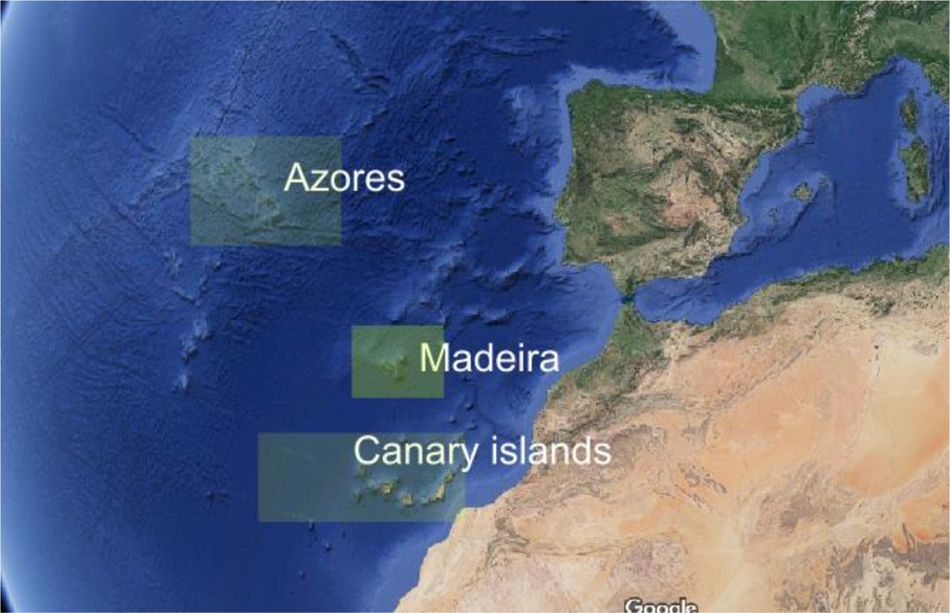
Map showing the distribution of the Macaronesian laurel forest in the Atlantic.

Several methods have so far been used to assess foliage susceptibility of potential hosts to *Pr* simulating optimal environmental conditions. The zoospore point inoculation on detached leaves is extensively used with *Phytophthora infestans* on potato leaves to select cultivar resistance or to determine and quantify key epidemiological parameters, such as latent period, leaf necrotic areas and sporangial density (Flier et al., 2003). Dipping excised host organs or whole plants in a zoospore suspension has been conducted for estimating the foliage susceptibility to *Pr* of tree forest species from the UK (Denman et al., 2005), potential ornamental plant hosts (Tooley et al., 2004), and forest trees (Hansen et al., 2005; Harris et al., 2020). Sporulation potential among plant species affected by SOD in California was assessed using leaf disc and leaf dip methods showing the last method produced two to four times lower sporangial counts (Rosenthal, Fajardo & Rizzo 2022). Although all these inoculation tests have predicted *Pr* natural infections relatively well, they tend to overestimate disease risk because they ignore some biotic and environmental factors, such as host phenology and genetic variation, temperature, inoculum density, and the plant’s architecture, which determine disease outcomes in the field (Hansen et al., 2005; Garbelotto et al., 2020). Nonetheless, these methods accurately assess the host response to infection and colonization under conditions suitable for the pathogen. Taking into consideration the advantages and flaws of the different inoculation methods, we aimed to test the foliage susceptibility of MLF species to infection in the laboratory and the capacity to sustain *Pr* sporulation as an approach for assessing the disease risk posed by *Pr* to this unique ecosystem. The MLF, an ecosystem ranging in species diversity between the oak-laurel forests of Asia and California, provides a reasonable scenario for speculating on the disease ecology of the pathogen in its native habitat.

## Material and Methods

### Isolates and inoculum production

Five isolates of *P. ramorum* were used in the experiments, three from the European lineage EU1, P1376 (obtained from *Viburnum tinus* cv. Eve Price), P1577 (from *Rhododendron catawbiensis*) and P1578 (from *R. grandiflora*), and two from the North American clonal lineage NA1, P1403 (*Vaccinium ovatum*; Oregon, USA) and P1579 (*Quercus agrifolia*, CA, USA). Stock cultures were maintained on carrot agar (CA) at 20° C in darkness (Moralejo et al., 2009). For the inoculation experiments, isolates were grown on CA in 90 mm diam. Petri dishes for 10-14 days at 20°C under continuous white light. These conditions promote the formation of large numbers of deciduous sporangia in culture. To obtain a sporangial suspension, *ca*. 10 ml of sterile distilled water (SDW) was added to each plate and the lids were sealed with parafilm. The sporangia were dislodged by vortexing the plates for 10 s at 40 hertz (Velp, Scientifica). The resulting suspension was decanted into an unused sterile Petri dish. Zoospore release was induced by chilling the dishes at 4°C for one hour before being returned to room temperature for 30 min. The concentration of the zoospore suspension was adjusted to 2−4 x 10^5^ zoospores per mL based on haemocytometer counts. For wound inoculations (explained below), mycelial plugs were obtained with a sterilised cork-borer 7 mm in diameter from the margins of 7 to 9-day-old cultures grown on CA.

### Plant material

Leaves of Macaronesian laurel forest species were collected from the Botanical Garden of Soller in Mallorca (Spain) on the day of the inoculation experiments. The species tested are listed in Table 1. Over 50 healthy, fully expanded leaves randomly selected from 2-4 individuals of each potential host species were collected early in the morning and put in plastic bags for transporting immediately to the laboratory. Here, the leaves were surface-disinfected with 70% ethanol for 10 s, rinsed in sterile distilled water and air-dried in a laminar flow hood. For each species, sets of four to six leaves, depending on their size, were selected randomly and placed with the adaxial face down on a metal grid in moist chambers consisting of transparent plastic boxes lined with sterile paper towels soaked in water.

**Table 1.** Plant species of the laurel forest of the Macaronesian region used in the inoculation experiments.

| Species | Family | Life form |
| --- | --- | --- |
| *Persea indica* | Lauraceae | evergreen tree |
| *Ocotea foetens* | Lauraceae | evergreen tree |
| *Laurus canariensis* | Lauraceae | evergreen tree |
| *Apollonias barbujana* | Lauraceae | evergreen tree |
| *Arbutus canariensis* | Ericaceae | evergreen tree |
| *Erica arborea* * | Ericaceae | heath tree |
| *Ilex canariensis* | Aquifoliaceae | evergreen tree |
| *Ilex perado* | Aquifoliaceae | evergreen tall shrub |
| *Viburnum tinus* subsp. *rigidum* | Caprifoliaceae | evergreen shrub |
| *Sambucus palmensis* | Caprifoliaceae | deciduous tree |
| *Myrica faya* | Myricaceae | evergreen shrub |
| *Picconia excelsa* | Oleaceae | evergreen tree |
| *Pistacia atlantica* | Anacardiaceae | evergreen shrub |
| *Visnea mocanera* | Theaceae | evergreen tree |
| *Salix canariensis* | Salicaceae | deciduous tree |
| *Heberdenia excelsa* | Myrsinaceae | evergreen tree |

\* Because the linear leaves of *Erica arborea*, a different inoculation method was performed and data of this species were excluded in the subsequent analysis. Other species usually found in this laurel forest include *Rhamnus glandulosa*, *Erica scoparia*, *Prunus lusitanica* and *Castanea sativa*.

### Inoculation techniques

We carried out inoculations on both wounded and unwounded leaves of 15 woody species (Figure 2A-F). In the first set of experiments, from April to May 2005, leaves were wound-inoculated with mycelial plugs. In the second set, done in the spring (April) of 2006, healthy intact leaves were inoculated with zoospores.

**Figure 2.**
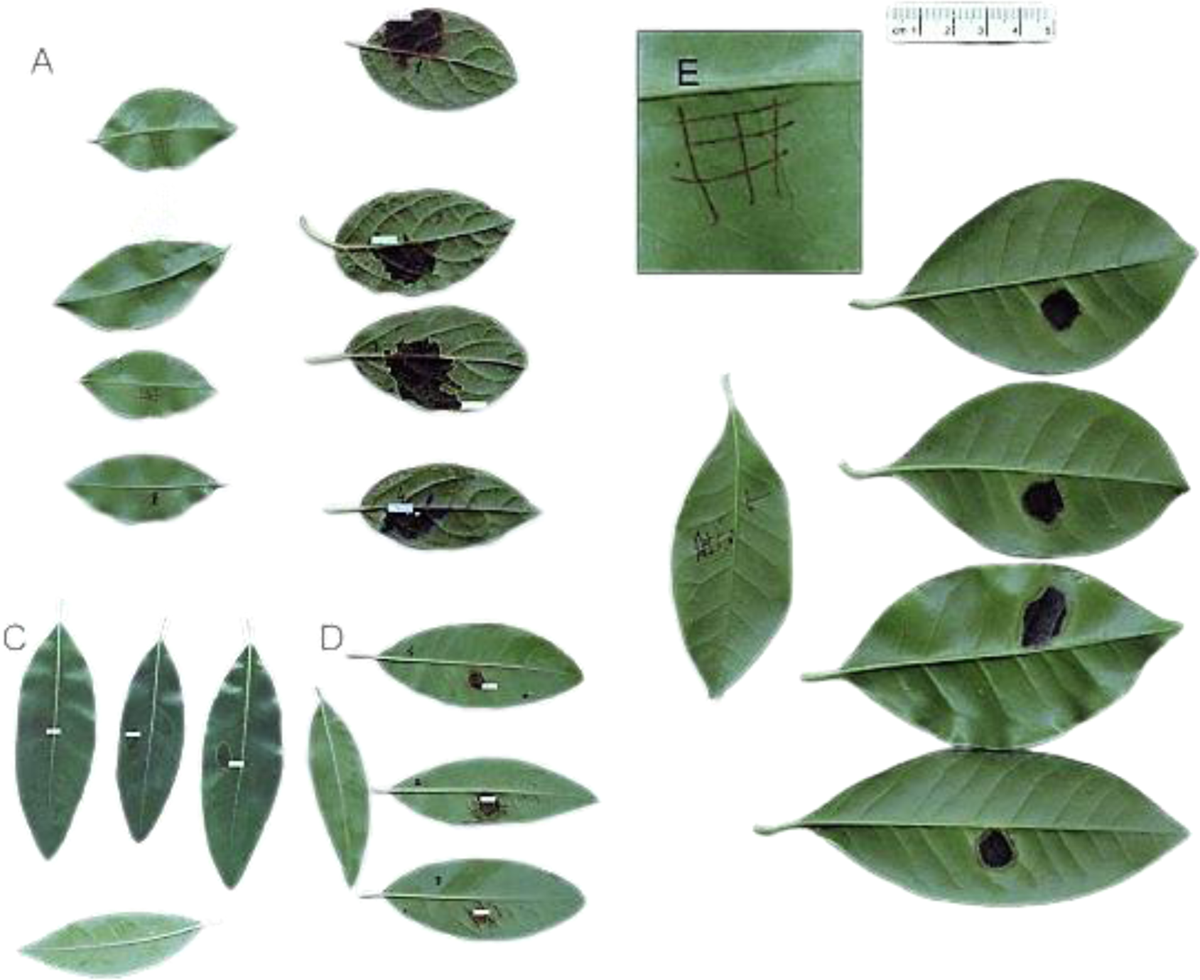
Leaf responses of Macaronesian laurel forest species to different inoculation treatments with *Phytophthora ramorum*. (A) Resistance of *Visnea mocanera* to isolate P1578 after 7 days (wound inoculation). (B) Susceptibility of *Viburnum tinus* to P1403 (zoospore-unwounded). Symptoms on the abaxial (C) and adaxial (D) leaf surface of *Persea indica* inoculated with P1376 four days after inoculating with zoospores. (E) Detail of the wounding procedure. (F) Tolerance response *of Picconia excelsa* to P1403 (wound inoculation).

#### Wound inoculation with mycelial plugs

Prior to inoculation, an area of approximately 1 cm^2^ of the lower surface of 4-6 leaves was wounded by lightly cutting the leaf in perpendicular strokes (a 3 x 3 grid) with a heat-sterilized razor blade (Figure 2E). The leaves were then inoculated by inverting a 6 mm diam. colony plug over the scraped surface. One of the leaves was used as a negative control by placing a sterile 6 mm diam. agar plug. 100 μl drops of SDW were placed over the plugs to maintain humidity. The moist chambers were incubated at 20°C and exposed to a 12 h photoperiod over seven days. Sporangial production was not measured in these tests.

#### Point (unwounded) inoculation with zoospores

Two separate tests were carried out because the evaluation required destructive sampling. Both lesion area and sporangial production were assessed on the fourth and seventh days after inoculation. in separate moist chambers, as independent assays. Ten moist chambers were thus prepared for each plant species tested (2 measurement dates x 5 isolates). The abaxial side of the leaf was inoculated by placing a single 100 μl drop of zoospore suspension near the centre of the midrib. One of the leaves was used as a negative control by placing a drop of sterile-distilled water on the leaf. All the boxes were incubated at 20°C under cool white light from fluorescent tubes suspended 30 cm above the chambers (12 h photoperiod). Four and seven days after inoculation, after removing sporangia for counting (see below), the leaves were scanned and the lesion area was calculated with the Olympus DP12 Soft version 3.2 (Figure 2B-D).

### Sporangial counting

For assessing sporangial formation on detached leaves we used the same inoculated leaves as in the above experiment. Sporangial induction followed S. Denman (http://rapra.csl.gov.uk/protocols) but is described briefly here. On day 3 for the 4-day incubation and on day 6 for the 7-day incubation, a single 100 μL drop of SDW was placed on the original inoculation point and leaves were further incubated under continuous white light for 24 h (as above). The water droplet was then suctioned off the leaf with a micropipette and transferred to a 1.5 ml centrifuge microtube. The leaf surface below the water drop (including all the necrotic parts resulting from the infection) was then gently scraped with a flamed spatula to dislodge sporangia. A 200 μl drop of sterile distilled water was placed on the lesion, the scrapings suspended in the drop and then collected in a 1.5 ml centrifuge microtube containing a drop of cotton blue in lactic acid. The centrifuge microtube was vortexed at high speed for 30 s, and five 10 μl aliquots of the suspension pipetted onto a microscope slide.

Total sporangial production and sporangial density (the number of sporangia per cm^2^) of each inoculated leaf were calculated. When ending the experiments, we became aware that *Pr* could asymptomatically infect leaves in some species and sporulate on them; therefore, we checked on which plant species this happened.

### Data interpretation and analysis

Laurel forest species responses to the different treatments were grouped as (i) symptomatic infection when conspicuous necrotic lesions developed; (ii) asymptomatic infection when no visible necrosis developed but sporangia were produced and the pathogen was re-isolated from the host; and (iii) no infection, when the pathogen could not be re-isolated. In the symptomatic response, two kinds of reactions were observed: compatible reaction with delayed host response, i.e. without significant difference between the fourth and seventh days after inoculation; and compatible reaction with reduced host response, i.e. lesion area significantly higher at day seven than at day four in the zoospore inoculation. To test infectivity, two pieces of *ca*. 5 x 5 mm tissue of the inoculated leaf were removed and plated on a semi-selective PARP medium (Erwin & Ribeiro 1996). Whenever a colony of *Pr* grew, the infection was considered positive. The infection efficiency (IE), measured as the percentage of leaves showing lesions out of the total number of leaves inoculated per species, was recorded.

All statistical analyses were performed using the *R* environment (*R* Development Core Team 2019). To know whether *Pr* isolates differed in their virulence to foliage and in their capacity to sustain sporangial production across MLF species, we used a generalized linear mixed-effects model (GLMM), treating tree *species* as a blocking random factor, *wounding* and *day* as a fixed factor and *lesion area* and *sporangia* as the response variable. The GLMM model for sporangia production only included *day* and *isolate* and their interaction as fixed categorical predictor variables, whereas those for *lesion area* also included the effect of the *wounding* treatment and its interaction with the *day*. Species that did not sustain sporulation were excluded from the analysis in the sporangia model. Controls were also excluded as lesions and sporangia did not develop on them. The time required to reach the sporulation peak from infection in polycyclic diseases is crucial to develop predictive epidemiological models of diseases caused by aerial Phytopthoras (Gómez-Gallego et al., 2019). To test whether the sporangial number increased between the fourth and seventh day of inoculation, we fitted a negative-binomial mixed model (estimated by ML and Nelder-Mead optimizer) using the function glmer in the R package lme4 (Bates et al., 2018), with *day* as a fixed factor, *day* x *isolate* interaction and *species* as random effects. To show differences in *sporangia* production among MLF species, we built a ridgeline chart as a density plot using ggplot2 and ggridges packages. Differences in *lesion areas* between the wounding and non-wounding treatments and between the 4^th^ and 7th-day assessments in the zoospore inoculation were modelled using *isolate* nested within *species* as random factors. To determine if there were differences between sporulation at 4 and 7 days after inoculation, we performed a mixed linear model analysis with MLF tree *species* as a random blocking factor and *isolates* as a fixed grouping factor. To determine if there was any relationship between lesion areas and sporangial density, we performed a GLMM using nle function with tree *species* as a fixed grouping factor and *lesion area* as the continuous independent variable.

### Climatic suitability

SOD epidemics in California and sudden larch death in the UK are driven by a concatenation of rainy episodes within temperature ranges optimal for sporulation (Meentmeyer et al., 2011; Harris et al., XXX). To compare the climatic suitability in areas of MLF with areas where Pr is established on the west coast of the USA, the UK, Ireland and France, North Vietnam and Japan, we used a climatic suitability index, *I*_t_ = *m_it_c_it_*, based on the weather conditions of Meentemeyer et al. (2011) model with few modifications. A moisture suitability index, *m_it_*, between 0 and 1, was calculated as the number of days in week t with precipitation greater than 2.5 mm in cell i. For the temperature suitability index, *c_it_*, instead of using the data of the mean production of zoospore at different temperatures (Davidson et al., 2005), we fitted a Beta function to data of sporangia production as a function of temperature (Englander, Browning & Tooley, 2004) (Fig. S1). This Beta function was used to convert the weekly average temperature based on daily temperature data in cell *i* during week *t* to the cell’s suitability index *c_it_*. This climatic suitability index expresses the average number of optimal annual infective-dispersive cycles of *Pr* for the period 1980-2020 without taking hosts into account.

Georeferenced data on the distribution of SOD in California were taken from the California Oak Mortality Task Force (https://www.suddenoakdeath.org/about-california-oak-mortality-task-force/), from areas with diseased larch from the Forest Research (https://data-forestry.opendata.arcgis.com/) and the areas of Vietnam and Japan from Jung et al. (2020). Worldwide temperature data were downloaded from the ERA5-Land dataset with hourly temporal resolution and 0.1° spatial resolution using GRIB format. To compute the annual Climatic suitability index a Julia library was built on top of GRIB.jl package.

## Results

### Lesion area

Lesion areas and infection efficiencies (IEs) were determined for detached wounded and unwounded leaves of 15 MLF species (Table 2). A total of 885 inoculations were performed. Among these, 600 (67.8%) induced necrosis, 124 (14.0%) leaves were asymptomatically infected (sustaining sporulation) and 161 (18.2%) remained uninfected, i.e. with the absence of symptoms and the pathogen could not be re-isolated. MLF species differed significantly in their susceptibility to *Pr* in the wounded (GLMM: χ2 =760.5, df=14; *p* <0.001) and unwounded treatments (on the 4^th^ day, GLMM: χ2 =499.0, df=10; *p* <0.001; and on the 7^th^ day, GLMM: χ2 =567.8, df=10; *p* <0.001). Eleven of the 15 laurel forest species inoculated developed necrotic lesions to different extents through zoospore inoculation, while they increased to 13 species through wound treatment (Table 2). Overall, 86.6% of the tested species could in some manner be considered susceptible to *Pr* when wounded. As an indicator of potential disease incidence at the plant community level, seven of the susceptible species had an IE of over 97% when kept unwounded and inoculated with zoospores (Table 2).

**Table 2.** Foliage susceptibility of laurel forest species from the Canary Islands to *Phytophthora ramorum*.

| Species | IE<sup>a</sup> (%) | Lesion area<sup>b</sup> (cm<sup>2</sup>) |  | Sporangia<sup>c</sup> |  |
| --- | --- | --- | --- | --- | --- |
|  |  | 4d | 7d | 4d | 7d |
| *Apollonias* | 0 | 0 | 0 | - | - |
| *Arbutus canariensis* | 100 | 1.9 (1.4-3.3) | 4.2 (2.9-5.3) | 91 (0-243) | 52 (12-126) |
| *Heberdenia excelsa* | 0 | 0 | 0 | - | - |
| *Ilex canariensis* | 97.5 | 1.8 (0.6-3.7) | 2.6 (0 -4.7) | 311 (37-922) | 101 (7-371) |
| *Ilex perada* | 10 | 0.15 (0 -0.9) | 0 | 47 (0-117) | 49 (11-139) |
| *Laurus azorica* | 98 | 0.85 (0.45-1.25) | 0.9 (0.3-1.6) | 80 (35-192) | 59 (20-209) |
| *Myrica faya* | 70 | 0.64 (0-1.3) | 0.54 (0-1.2) | 73 (33-178) | 75 (13-185) |
| *Ocothea foethens* | 50 | 0.27 (0-0.84) | 0.33(0-0.96) | 48 (4-98) | 72 (8-352) |
| *Persea indica* | 100 | 1.0 (0.5-1.5) | 1.1 (0.71-1.84) | 72 (9-164) | 146 (27-382) |
| *Picconia excelsa* | 0 | 0 | 0 | 113 (0-286) | 164 (13-690) |
| *Pistacia atlantica* | 100 | 1.04 (0.54-1.54) | 1.02 (0.66-1.37) | 201(72-446) | 124 (8-296) |
| *Salix canariensis* | 100 | 0.5 (0.28-0.77) | 0.55 (0.3-1.1) | 95 (33-218) | 95 (18-384) |
| *Sambucus* | 32.5 | 0.65 (0-2.4) | 0.48 (0-3.97) | 25 (0-90) | 48 (0-193) |
| *Viburnum tinus* | 100 | 4.75 (1.9-10.1) | 8.12 (2.48-19.0) | - | 200 (57-578) |
| *Visnea mocanera* | 0 | 0 | 0 | - | - |

The infection efficiency (IE), lesion area and capacity to sustain sporulation of *P. ramorum* were assessed on detached leaves inoculated with a 100 µL droplet of a zoospore suspension on the leaf underside. Four to six leaves of each species were inoculated per each isolate used (three European (EU1) and two North American lineages).
<sup>a</sup> Infection efficiency: the percentage of inoculated leaves that developed necrotic lesions (*n* = 30-60).
<sup>b</sup> mean lesion area (and range of scores).
<sup>c</sup> mean number of sporangia per leaf (and range of scores)

We found significant differences in lesion area on the 7th day after inoculation between *Pr* isolates across laurel forest species for both wound inoculation (GLMM: χ2 =17.7, df=4; *p* = 0.0014) and unwounded inoculations (GLMM: χ2 =10.97, df=4; *p* = 0.026). However, there were no differences in the unwound inoculation at the 4^th^ day (GLMM: χ2 =1.93, df=4; *p* = 0.75). The three EU lineage isolates were on average 8.6% (mean =1.9 ± 1.9) more virulent than the two American isolates (mean =1.75 ± 2.0) on the 7^th^ day after inoculation, which agrees with other inoculation studies using the same isolates (Moralejo et al., 2009) or clonal lineages (Manter et al., 2010). Wounding substantially increased the IE on all plant species versus zoospore inoculations on intact leaves (Table 2). However, the effect of wounding on lesion expansion varied among hosts. For example, in *Arbutus canariensis*, *Ilex canariensis*, *Persea indica* and *Pistacia atlantica* lesion areas were significantly larger in the unwounded than in the wounded inoculations, whereas in *Sambucus palmensis*, *Myrica faya*, *Ocotea foetens*, *Picconia excelsa*, *Heberdenia excelsa*, *Ilex perado*, *Apollonias barbujana* and *Visnea mocanera* wounding significantly increased lesion areas (Figure 3).

**Figure 3.**
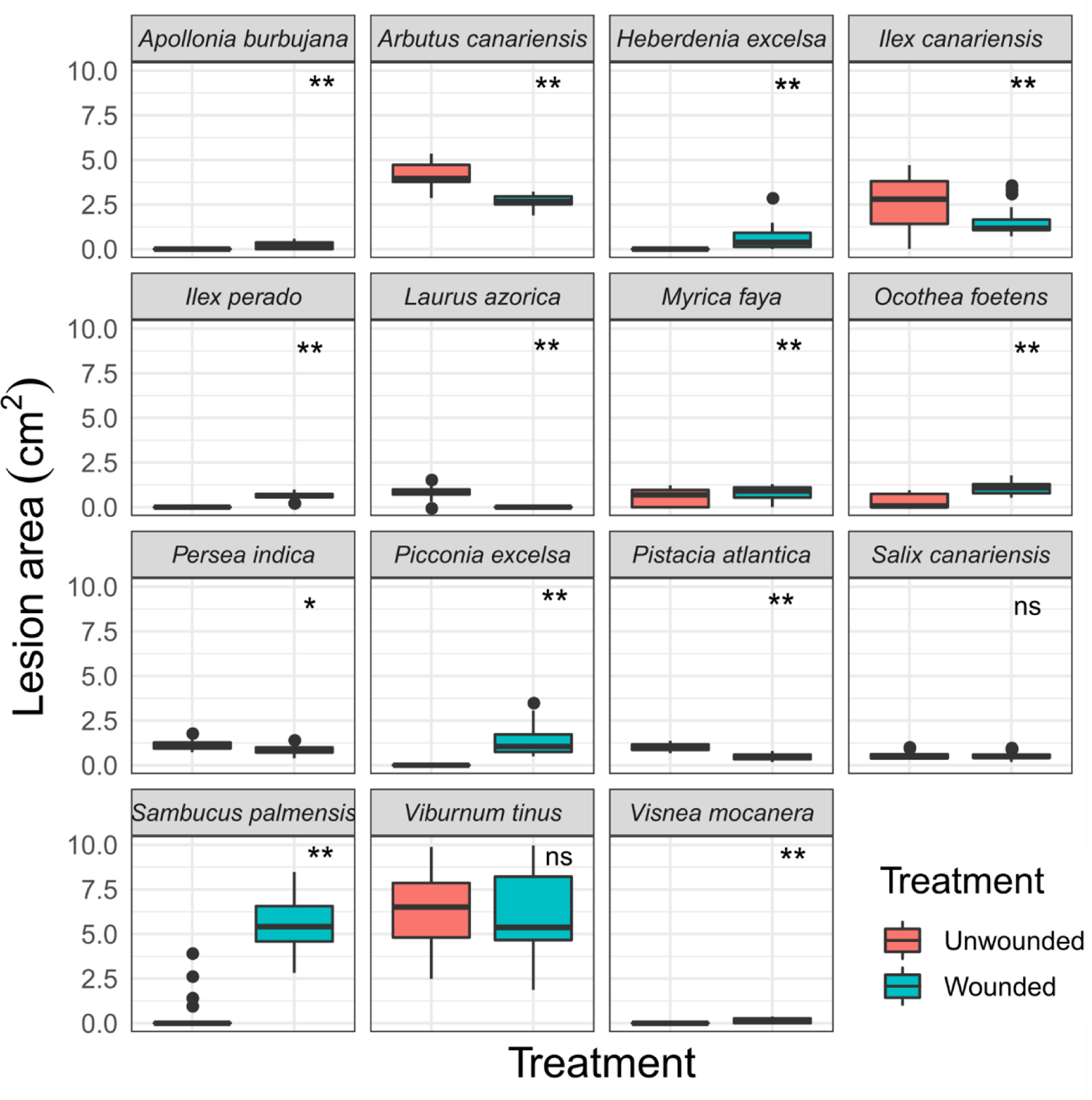
Boxplot showing lesion areas caused by *Phytophthora ramorum* on unwounded and wounded leaves of MLF species seven days after inoculation. Pair-contrast tests between unwounded and wounded treatment were performed for each plant species; ns = non-significant, * significant (*p* < 0.05), ** highly significant (*p* < 0.001).

Leaves of *P. excelsa* and *H. excelsa* only developed necrotic lesions after wound treatment. However, those of *P. excelsa* sustained *Pr* sporulation, which indicates that there was an infection. In the case of *H. excelsa,* a physical barrier may have impeded penetration after inoculation. By comparison, lesions were up to 3.0 cm^2^ when leaves were wounded. Two species, *A. barbujana* and *V. mocanera*, formed relatively small lesions in the wounded area but none after zoospore inoculations.

Other species could not be assigned to a susceptible class; *Ilex perado* and *Ocotea foetens* showed a significantly lower IE after zoospore inoculation compared to wound treatment (Table 2). In *Myrica faya*, *Laurus azorica*, *Persea indica*, *Pistacia atlantica* and *Salix canariensis*, IE was *ca*. 100% after unwounded inoculation but lesion expansion differed only slightly on the fourth and seventh days after inoculation, lesion areas seldom exceeding 2 cm^2^ (Figure 4). By contrast, in three other species, *Arbutus canariensis*, *Ilex canariensis* and *Viburnum tinus*, there were significant differences (*P* < 0.001) in lesion areas between the fourth and seventh days of incubation (Figure 4). The most susceptible host was *V. tinus* (Caprifoliaceae), where lesions up to 19 cm^2^ were recorded seven days after inoculation, followed by *S. palmensis* (Caprifoliaceae) (mean = 6.3 cm^2^) when wounded.

**Figure 4.**
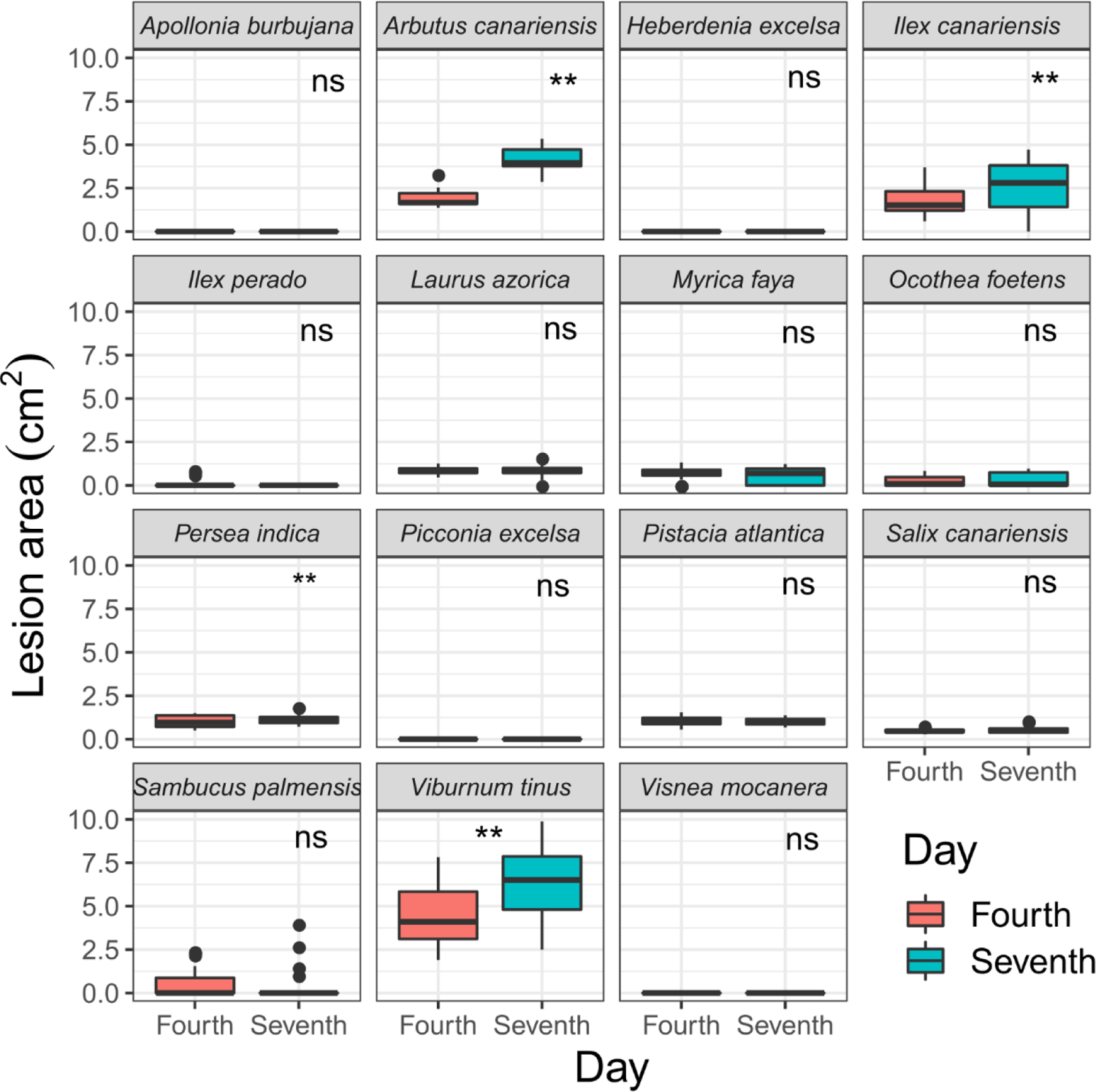
Boxplot showing lesion area formed by *Phytophthora ramorum* on leaves of species from the Macaronesian laurel forests four and seven days after inoculating with zoospores. Pair-contrast *t*-tests between the 4th and 7th day tests were performed for each plant species; ns = non-significant, * significant (*P* < 0.05), ** highly significant (*P* < 0.001).

### Sporangial production

The leaves of most plant species inoculated, even those that did not develop necrosis, sustained *Pr* sporulation (Table 2 & Figure 5). Levels of sporangial production on leaves varied significantly between laurel forest species on the 4^th^ day after inoculation (GLMM: χ2 =214,9, df=11; *p* <0.001) and on the 7^th^ (GLMM: χ2 =83.7, df=11; *p* <0.001). *Ilex canariensis* and *Pistacia atlantica* sustained the highest level of sporulation between species on the 4^th^ day, whereas *Viburnum tinus* and *Picconia excelsa* on the 7th day (Wald-z, *p* < 0.05). Strikingly, when counted across all plant species, sporangial production per leaf was not significantly higher on the seventh than on the fourth day after inoculation (GLMM: χ2 = 0.87, df=1; *p* = 0.35), with sporangia production 15% higher at the 4^th^ day. The rate of sporangial production was significant when measured as sporangia per lesion area (GLMM: χ2 = 10.71, df=1; *p* = 0.001) and was 41% higher on the fourth day. Slopes of linear regressions between sporangia production and lesion area showed negative trade-offs across species (Figure 5). The latent period, defined as the time from inoculation to the first evidence of sporulation, was less than 4 days in all species tested and similar to that of other aerial dispersed species, such as *Phytophthora infestans*.

**Figure 5.**
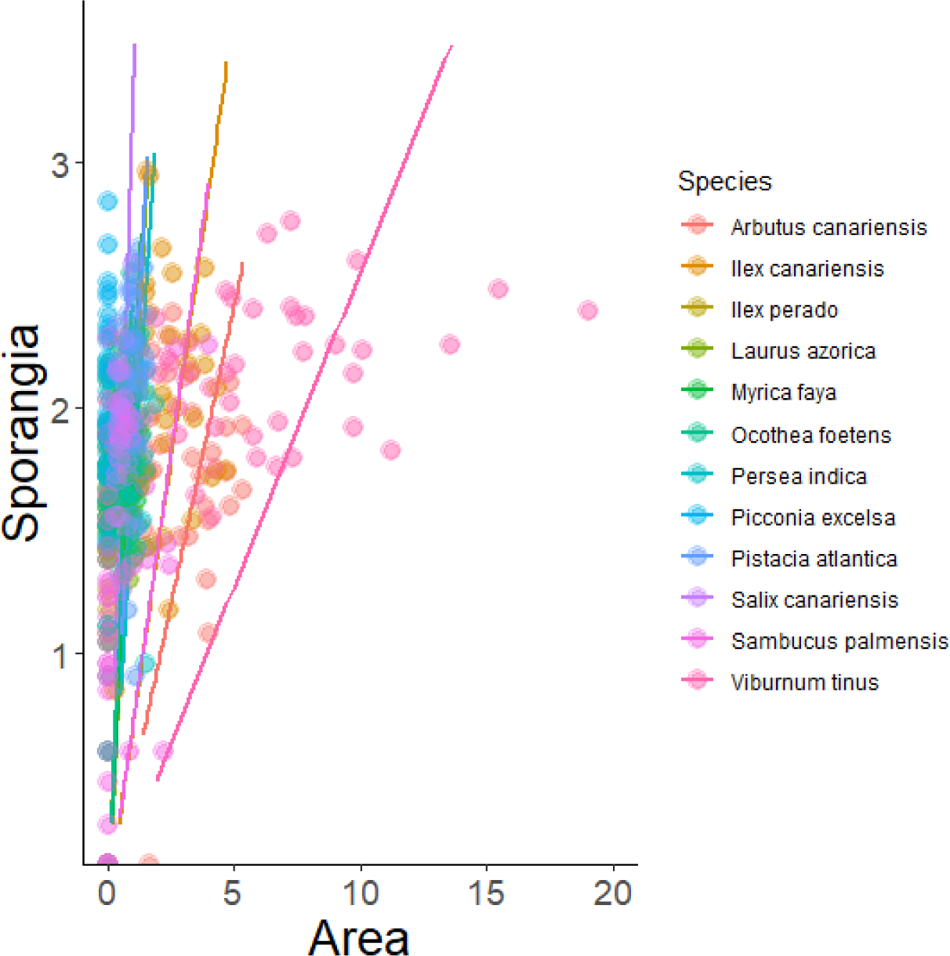
Scatter plot of the lesion area caused by *Phytophthora ramorum* on leaves of 15 Macaronesian laurel forest (MLF) species against the sporangia number after inoculation on unwounded detached leaves. The slopes suggest negative trade-offs between pathogenicity and sporulation among MLF species.

**Figure 6.**
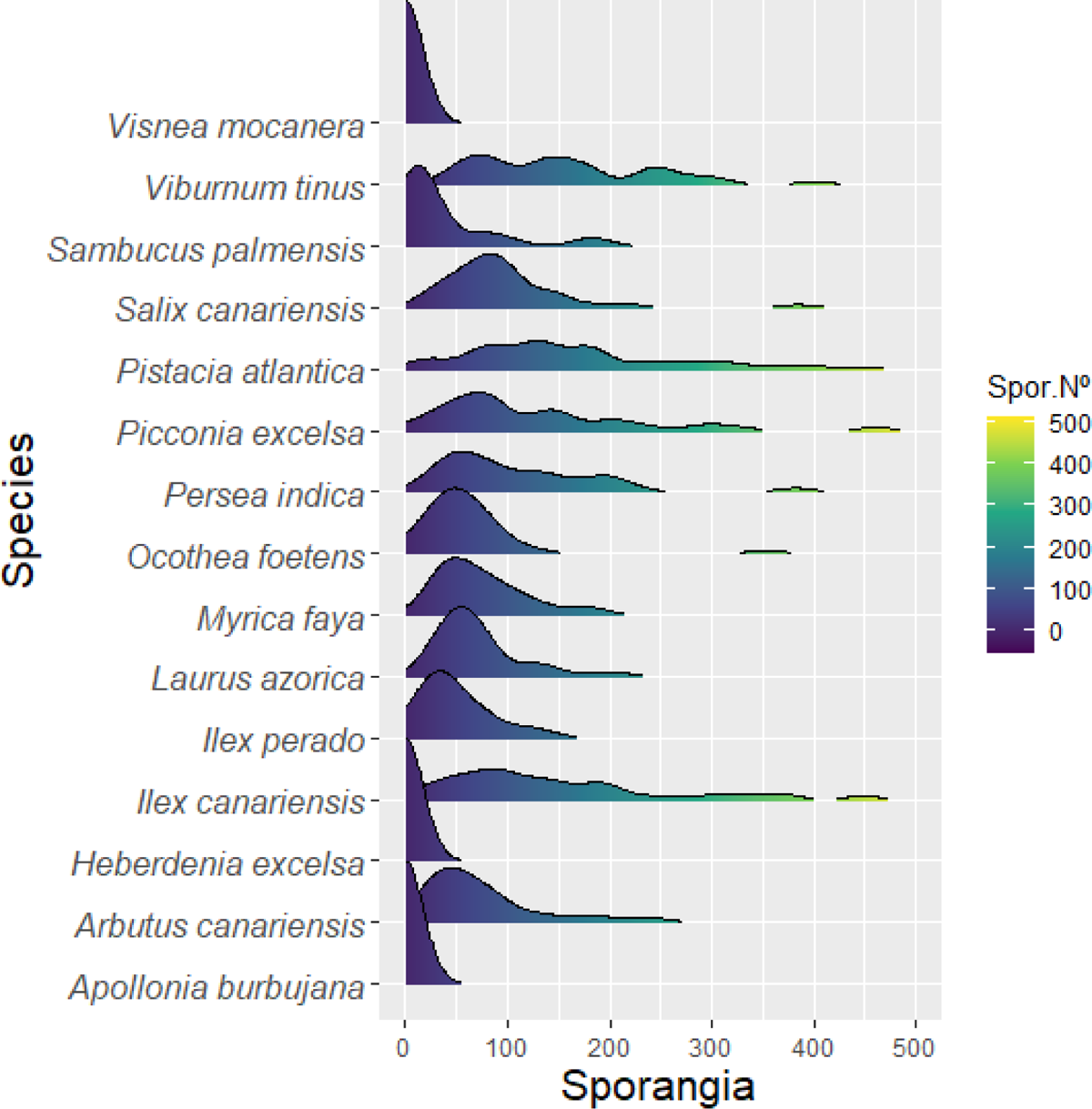
*Phytophthora ramorum* sporangial production across MLF species. A ridgeline plot showing the distribution of sporangial production of *P. ramorum* on leaves of MLF species *in vitro*.

### Climatic suitability

We found relatively low climatic suitability indices for *Pr* in MLF areas compared to laurel forests in northern Vietnam and Japan, but similar to areas with SOD in California (Tukey post-hoc test, *p*=0.94), as expected given that both areas share a Mediterranean climate (Fig. 7).

**Figure 7.**
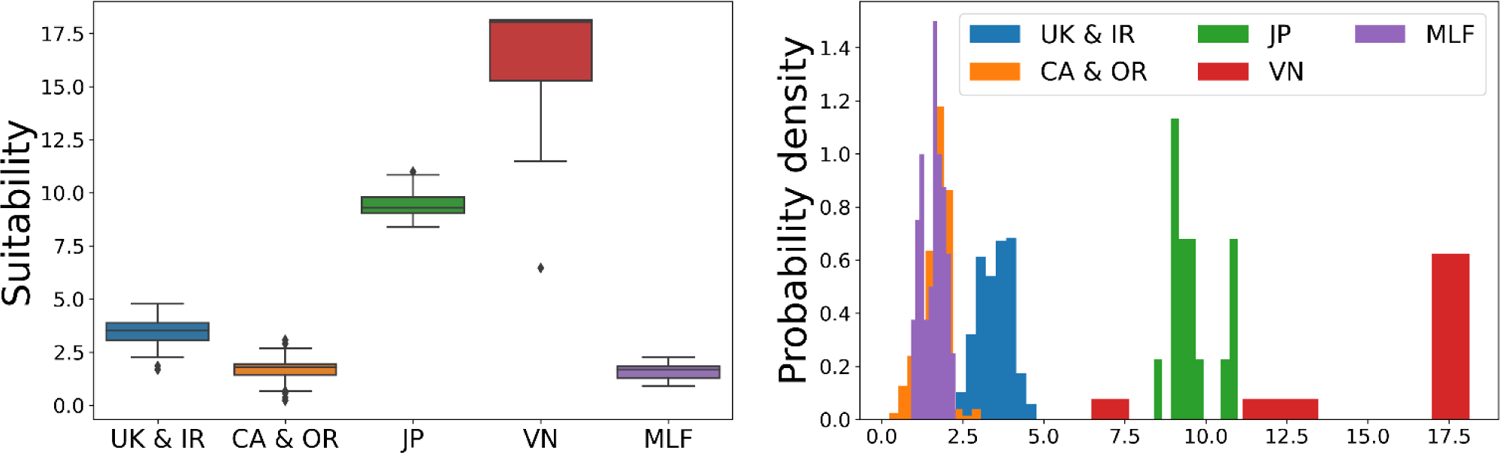
Climatic suitability of Macaronesian laurel forest (MLF) in relation to areas where *Phytophthora ramorum* is established. (a) Boxplot of suitability index which reflects the average number of potential sporulation-dispersal events from 1980 to 2020. (b) Probability density of suitability index for MLF and UK + IR = United Kingdom + Ireland; CA + OR = California + Oregon; JP = Japan; VN = Vietnam. Suitability indexes among zones differ significantly (*F*_4,1024_ = 2523, p < 0.001)

These low average climatic suitability index values, however, are consistent with field studies of seasonal sporangia production in SOD-affected areas in California (Davidson et al., 2011), with observed substantial increases in sporangia in exceptionally wet springs during El Niño periods (Davidson et al., 2005), and with greater sporangial captures and lesion survival on *U. californica* leaves in the wetter redwood forest than in mixed-the evergreen oak forest.

## Discussion

In this study, we show how five isolates of *Pr* with diverse origins can infect to the same degree detached leaves of the majority of woody plants of a moderate species diversity ecosystem such as the MLF, where humid to hyper-humid conditions prevail throughout the year. Similar results of a potential host range (> 70% of total species inoculated) have been reported for laboratory inoculations on species of European temperate forests (Denman et al., 2005; Harris & Webber, 2016), evergreen mixed forests of the USA Pacific coast (Hansen et al., 2005), temperate forests of the eastern USA (Tooley & Browning 2009), Australian flora (Ireland et al., 2012) and evergreen sclerophyllous woodlands in the Mediterranean (Moralejo et al., 2006b; Moralejo et al., 2009). Such infection capacity suggests adaptations of the pathogen (multiple-host strategy) in their ecosystems of origin and the existence of common basal defence systems phylogenetically conserved in woody plants that *Pr* would be capable of overcoming (see Heath 1991; Moralejo et al., 2006a).

Of the three factors for disease development, a favourable environment, susceptible host(s) and presence of the pathogen, the MLF clearly meets the first two. Climatic parameters within MLF distribution, such as precipitation, mean T_max_ and T_min_, as well as mean relative humidity, fall in the range of those parameters collected in the ecological niche modelling for SOD distribution in California (Meentemeyer et al., 2008). SOD epidemics are primarily driven by the combination of winter and spring precipitation (av. 1251 mm), mild temperatures (av. T_max_= 17.2 °C; T_min_= 4.9°C), high relative humidity (69.2%) and the presence of bay laurel, whilst survival of inoculum in summer depends upon the additional moisture provided by fog (Rizzo et al., 2002). Likewise, winter precipitation in MLF ranges from 500 mm in the southern islands to higher than 1900 mm in some areas of Madeira and the Azores Islands; mean relative humidity is over 87%, and mean fog frequency in summer is over 89% (Marzol 2008) *vs*. 40-44% recorded in redwood and laurel-oak forest of central and northern California (Johnstone & Dawson 2010). Fossil records indicate that both redwoods and some MLF species coexisted in Europe and had a broader range under a warm and humid climate during the Tertiary (Axelrod 1975; Alcalde-Olivares et al., 2004). To date, both relict forests are restricted to highly fragmented and small distant coastal refugees that protected them from Plio-Pleistocene climate changes.

Thirteen of the 15 MLF species here tested were to some degree susceptible, of which seven had an IE of over 97% when inoculated with zoospores (Table 2). In other experiments (data not shown), the leaves of *Erica arborea*, an important component of MLF, were consistently susceptible to *Pr* when dipped in a zoospore suspension. Furthermore, other plant species commonly associated with the MLF, such as *Prunus lusitanica*, *Castanea sativa*, *Pittosporum undulatum* (naturalised invasive), *Erica scoparia* and *Rhamnus glandulosa*, have been reported as hosts (Denman et al., 2005; Hüberli et al., 2006) or belong to genera including many susceptible species.

To gain insights into the response of MLF species to infection, we performed inoculations on wounded and unwounded leaves taken during the same season and from the same plant individuals (i.e. genotypes). We observed a continuum of host responses from resistance, in some cases tentatively attributed to constitutive barriers, such as in *Visnea mocanera* (Figure 2A), to non-resistance (i.e. unrestricted pathogen colonisation), such as that exhibited by *Viburnum tinus* (Figure 2B). Compatible reactions in other eight MLF species also showed some kind of host defence responses, since their lesion areas did not differ, or differed only slightly between the 4^th^ and the 7^th^ day after inoculation (Figure 4). In these cases, sporulation on the lesion was not constrained, demonstrating some tolerance of MLF species towards the pathogen.

Two factors, pathogen’s transmission potential and host densities, mainly determine the likelihood of invasion, persistence and spread of infectious diseases within plant communities (Anderson & May 1986). Despite considerable research on SOD epidemiology, only until recently data on *Pr* sporulation on key foliage hosts were available for predicting disease risk in other forest ecosystems. In our experiments, the average sporangial production per laurel forest species (*n* = 15 taxa) on the seventh day was somewhat low (μ = 99 sporangia per leaf), but similar to those counted for bay laurel leaves (*Umbellularia californica*) in both laboratories (Hüberli et al., 2006; see Table 2) and natural infections in California (cf. Davidson et al., 2005; Davidson et al., 2008), as well as in laboratory trials in the UK (Sansford et al., 2009). The leaves of most of the MLF species tested by us, even those being asymptomatic, sustained relatively few sporangia (Table 2). Coincident with our results, Tooley & Browning (2009) found sporangial productions per leaf ranging from 2.5 to 198.2 on 25 inoculated native plant species from the eastern USA, and similar numbers have been counted on leaves of more than 25 inoculated Mediterranean plant species (Moralejo et al., unpublished). On the other hand, five species of the MLF, *Ilex canariensis*, *Persea indica, Pistacia atlantica, Picconia excelsa* and *Viburnum tinus* sustained sporangial numbers comparable to those found on bay laurel (Table 2). In addition, two species belonging to the Lauraceae, *Persea indica* and *Laurus azorica*, as is also the case of *U. californica*, showed similar IE values, lesion area and *in vitro* sporangial production per leaf in the case of *P. indica* to those reported for bay laurel (Table 2). We, therefore, believe that MLF would promote within- and between-host disease transmission if the pathogen becomes established.

Two selective pressures might determine the lifestyle of *Pr* in its original habitat(s): the constraint on short-distance dispersal in a moderate to high diversity species forest (e.g. laurel-oak forest) and the fitness costs caused by the need for a multiple-host strategy, which seems to be associated with reduced sporulation (Moralejo et al., 2006a). Unlike previous reports for bay laurel (Davidson et al., 2005), in our tests, the rate of sporangial production decreased over time in most MLF species as lesions expanded. This suggests a limited time for sporangial production and *Pr* transmission from existing lesions. Such low levels of sporangial production and short transmission periods combined with the short-distance dispersal (Davidson et al., 2005) require a high host density and frequent rain or mist events for *Pr* effective spread. The MLF fulfils these conditions, having a high mean leaf area index (LAI > 7; Morales et al., 1996), a potential multi-host canopy likely permitting similar rates of within- and between-host disease transmission, and a macro- and microclimate conducive to disease establishment and spread. On the basis of the results of our inoculations, the comparison of epidemiological parameters of *Pr* on its main host *U. californica* in California, the present knowledge of SOD ecology and the original habitat of *Pr* in oak-laurel forests, we predict a moderate to high risk of invasion of the MLF if the pathogen is introduced.

## Acknowledgements

The European Union, Project RAPRA 502672 (2004-2007), funded this research. Experiments were conducted under an officially authorised laboratory license (RE-9058/2003). We are very grateful to the Botanical Garden of Soller (Mallorca) for providing the plant material necessary for the experiments.

## Authors Contribution

José Andrés García-Muñoz & Eduardo Moralejo carried out the inoculation assays. Àlex Giménez-Romero developed the climatic suitability analysis and maps and Sandra Denman & Eduardo Moralejo analysed the data and wrote the manuscript.

## Supplementary Figures

**Supplementary figure 1.**
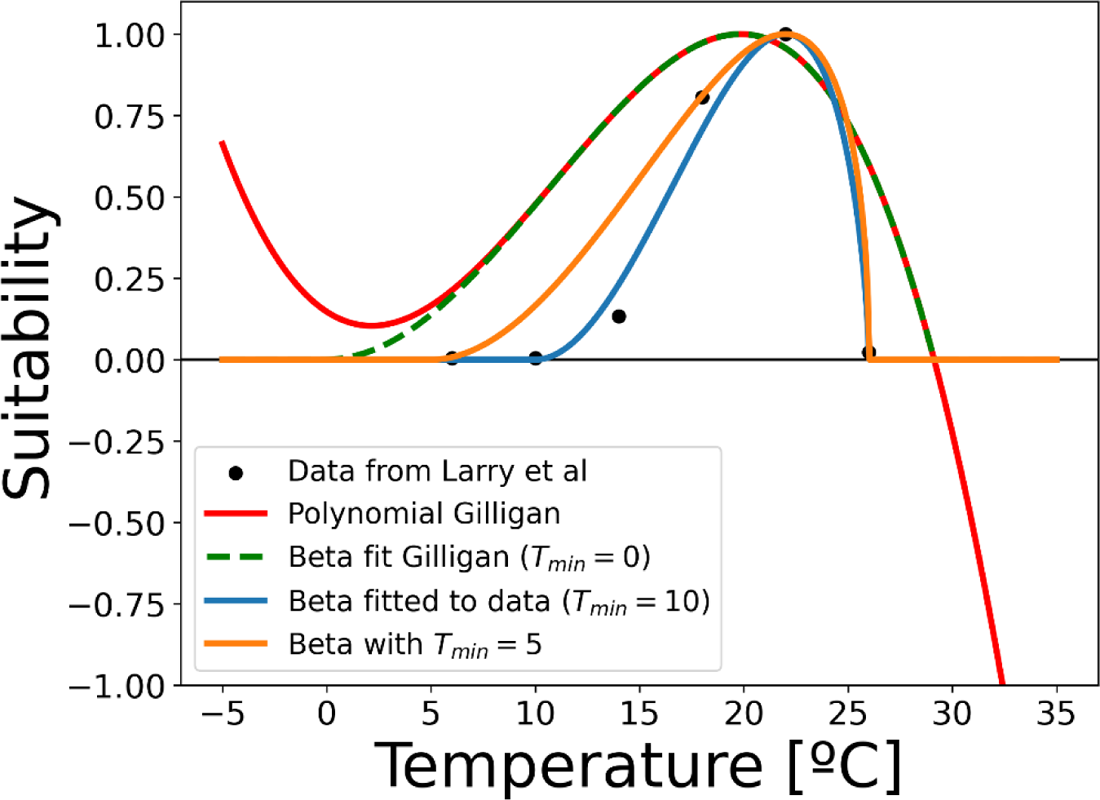
Relationship between the relative maximum sporulation of and temperature used in different models. We used the Beta with *T_min_* as it captures the infective-dispersal cycles at mean lower temperatures in more oceanic climates.

**Supplementary figure 2.**
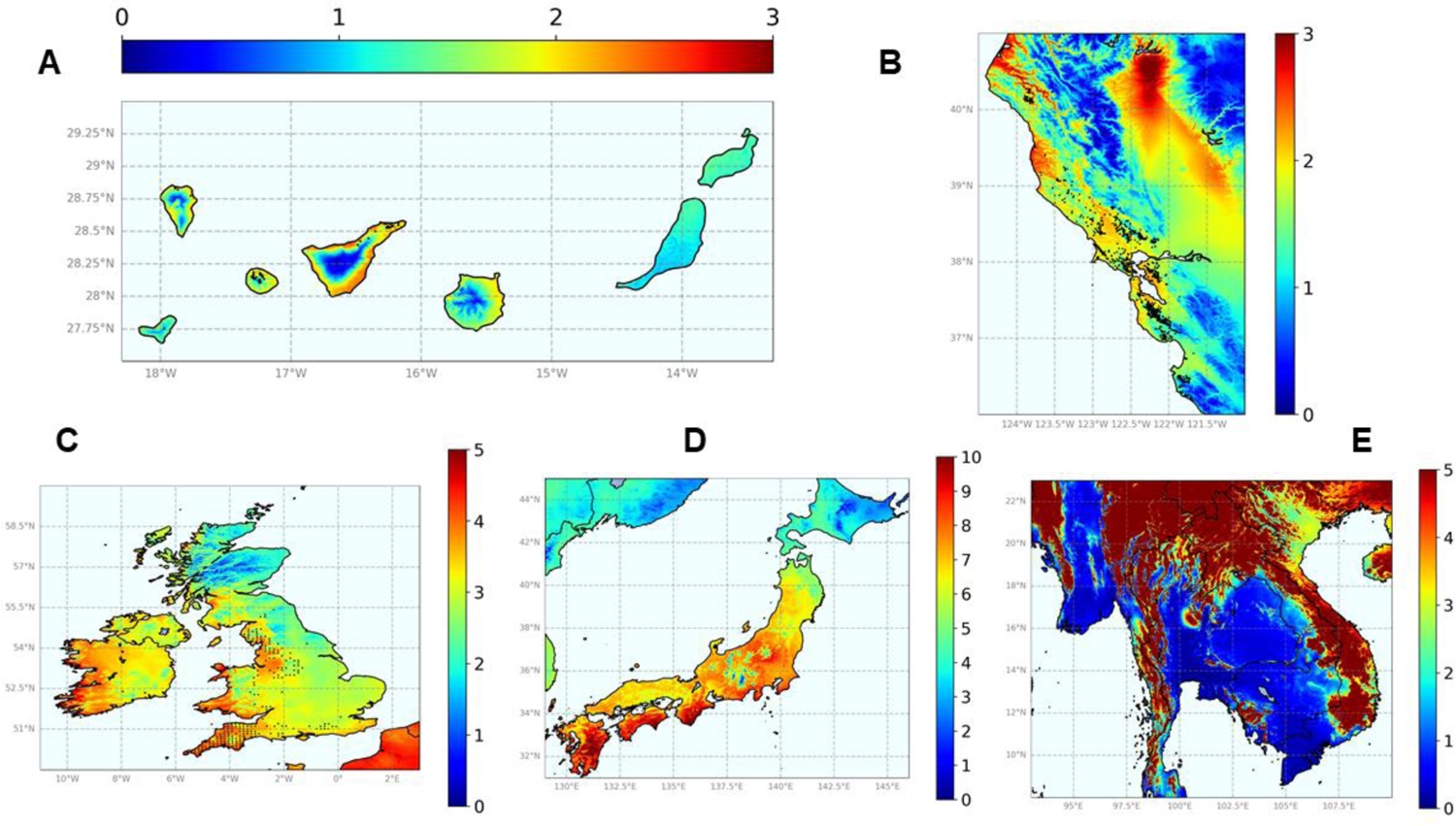
Maps showing the suitability index for *Phytophthora ramorum* establishment. The scales represent the average cumulative annual number of infectivity-dispersal events at optimal conditions during the period 2000-2019. These maps do not capture the climatic variability in which exceptional rainy episodes lead to large epidemics. (A) Canary Islands;(B) California and south Oregon;(C) UK and Ireland; (D) Japan; (E) South east Asia (Vietnam).

